# SCD1 inhibition synergizes with TMZ to improve IDH1^MUT^ glioma survival of mice via lipotoxic stress

**DOI:** 10.64898/2026.06.09.731222

**Authors:** Laurence Zhang, Tomohiro Yamasaki, Viney Kumar, Taylor A. Harmon, Helena Muley, Meili Zhang, Dionne Davis, Samarth Mathur, Ross Lake, Christel Herold Mende, Tyrone Dowdy, Adrian Lita, Mioara Larion

## Abstract

Mutations in the isocitrate dehydrogenase enzyme (IDH1) are prevalent in low-grade gliomas, such as oligodendrogliomas. Other than surgery, radiation, and chemotherapy, few options for treatment exist. The standard of care for patients currently consists of maximal safe resection, as well as the potential use of radiation and chemotherapy, typically followed by radiographic surveillance. Although significant advances have been achieved, these have not translated into meaningful improvements in overall survival, warranting the development of novel therapies. Herein, we demonstrate that the inhibition of SCD1 in vitro leads to decreased colony formation and is more specific to IDH1^MUT^ glioma than the IDH1^WT^ cells, We further identified MF-438 as an active inhibitor of this enzyme and showed that this inhibitor is linked with iron transport and ferroptosis. MF438 depleted oleic (C18:1) and palmitoleic acid (C16:1) levels, driving saturated phosphatidylcholine (PC) accumulation and ER stress (INSIG1, SEL1L upregulation). TS603 exhibited selective polyunsaturated phosphatidylcholine reduction with downregulation of GPX4, FTH1, and KEAP1, and upregulation of NCOA4, SLC11A2, ALDH7A1, and DPP4, consistent with ferroptosis priming via ferritinophagy-driven expansion of the labile iron pool, as confirmed by FerroOrange flow cytometry. Neutral lipid metabolism genes (LPIN1, LDLR, PNPLA3, ACSL1), intracellular lipid transport genes (TMEM41B, OSBP, STARD4), and lipid droplet organization genes (SQLE, CHKA, AUP1) were coordinately upregulated in TS603 after treatment with MF-438. MF438+TMZ activated the integrated stress response (ATF3, DDIT3, IRF1, CDKN1A, GADD45B) and synergistically suppressed TS603 neurosphere growth (p=0.0326). In vivo combination between MF-438 and TMZ showed improved survival versus the TMZ-alone group, in an IDH1-mutant oligodendroglioma model. Tissue analyses showed significant reduced Ki67 expression, a marker of cellular proliferation, in the combination treatment. Collectively, these findings suggest that targeting lipid metabolism may enhance the efficacy of standard-of-care therapy in IDH1-mutant oligodendroglioma.

## INTRODUCTION

Mutations in the isocitrate dehydrogenase enzyme (IDH1) are prevalent in low-grade gliomas, such as oligodendrogliomas^1–3^. Other than surgery, radiation, and chemotherapy, few options for treatment exist^2,4^ The standard of care for patients currently consists of maximal safe resection, as well as the potential use of radiation and chemotherapy with temozolomide, typically followed by radiographic surveillance^4^. For low-risk patients, treatment is less defined after surgery, often consisting of surveillance thus affecting their quality-of-life significantly^5^. Regardless of the risk level, these tumors eventually progress and generally are fatal. Thus, more effective and more personalized treatments are urgently needed.

Diffuse gliomas harboring mutations in isocitrate dehydrogenase 1 (IDH1) represent a biologically distinct subgroup of brain tumors characterized by extensive metabolic rewiring^23,6–9^. Through the production of the oncometabolite D-2-hydroxyglutarate (2-HG), mutant IDH1 alters epigenetic regulation^10–12^, cellular metabolism^9,10,13–17^, and redox homeostasis^8,13,18^, creating metabolic dependencies that distinguish these tumors from their IDH-wild-type counterparts. While these alterations have provided important insights into glioma biology, the metabolic vulnerabilities created by IDH mutations remain incompletely understood and may offer opportunities for therapeutic intervention.

Proliferating cells have a high demand for monounsaturated fatty acids (MUFAs), which are essential components of phospholipids, triacylglycerols, and cholesterol esters required for membrane biogenesis. Beyond their structural role, MUFAs also regulate cellular signaling pathways and influence interactions between tumor cells and the immune microenvironment. Consequently, many cancers exhibit enhanced lipid synthesis and remodeling to support growth, survival, and therapeutic resistance. Elevated MUFA levels have been associated with aggressive tumor behavior and poor clinical outcomes in multiple malignancies, including hepatocellular, thyroid, prostate, pancreatic, renal, melanoma, and breast cancers^19–25^.

The rate-limiting step in MUFA biosynthesis is catalyzed by stearoyl-CoA desaturase (SCD), which introduces a double bond into saturated fatty acids to generate MUFAs^26^. In humans, two SCD isoforms have been identified^26^. SCD1 is localized primarily to the endoplasmic reticulum (ER), where de novo lipid synthesis occurs, and is expressed in adipose tissue, liver, heart, lung, and brain. SCD5 is more restricted in its expression and is found predominantly in the brain and pancreas^27–29^. A growing body of evidence demonstrates that SCD1 is a critical regulator of cancer cell metabolism, promoting stemness, tumor progression, autophagy, metastasis, and resistance to therapy. Pharmacological inhibition of SCD1

reduces intracellular MUFA levels and produces potent antitumor effects in preclinical models, including glioblastoma^23,24^. Furthermore, studies of breast cancer brain metastasis have shown that SCD1 expression is required for efficient colonization of the brain, underscoring the importance of lipid desaturation pathways in the central nervous system tumor microenvironment^20^. Despite the well-established role of mutant IDH1 in metabolic reprogramming and the intimate connection between IDH1-driven metabolism and fatty acid biosynthesis^30^, the contribution of SCD1 to glioma biology, particularly in the context of IDH mutations, remains poorly defined.

We previously demonstrated that IDH-mutant glioma cells exhibit abnormal partitioning of MUFAs into endoplasmic reticulum (ER) and Golgi membranes, resulting in organelle dysfunction and increased susceptibility to lipid stress^16^. These findings raised the possibility that IDH-mutant glioma cells may be particularly dependent on SCD1-mediated lipid desaturation for survival. Because inhibition of SCD1 disrupts membrane homeostasis and promotes the accumulation of saturated fatty acids, targeting this pathway may expose a metabolic vulnerability unique to IDH-mutant tumors. In addition to its role in lipid metabolism, SCD1 activity is closely linked to cellular redox regulation. The desaturation process requires iron-dependent enzymes, suggesting a potential connection between iron homeostasis and lipid metabolic adaptation. However, the mechanisms coordinating these pathways remain largely unknown.

Disruption of lipid desaturation has also been linked to ferroptosis, a form of iron-dependent regulated cell death driven by lipid peroxidation^31^. Furthermore, severe metabolic stress and D-2HG in particular can activate additional regulated cell death pathways, including necroptosis^32^. These observations are particularly relevant in glioma, where resistance to conventional therapies remains a major barrier to durable treatment responses. TMZ, the current standard chemotherapeutic agent for oligodendroglioma and other diffuse gliomas, exerts its antitumor effects primarily through induction of DNA damage^33^. However, the efficacy of TMZ is frequently limited by adaptive stress-response pathways that allow tumor cells to survive treatment.

Given the central role of SCD1 in maintaining lipid homeostasis and the altered metabolic landscape of IDH-mutant gliomas, we hypothesized that inhibition of SCD1 would impair the ability of glioma cells to adapt to oxidative and metabolic stress, thereby sensitizing them to TMZ. In this study, we investigated the consequences of pharmacological and genetic SCD1 inhibition in IDH-mutant and IDH-wild-type glioma models. We identify a previously unrecognized functional relationship between SCD1 and DMT1, demonstrate that SCD1 inhibition profoundly alters lipid and iron metabolism, and show that these changes sensitize glioma cells to TMZ through activation of ferroptotic and necroptotic cell death pathways. Together, our findings establish a mechanistic link between lipid desaturation and iron homeostasis and highlight SCD1 as a promising therapeutic target for the treatment of IDH-mutant gliomas.

## METHODS

### Cell Culture

Glioma cell lines: TS603 and GSC923, were cultured with DMEM: F12 media (Gibco Laboratories, Gaithersburg, MD, USA) with the supplement of penicillin/streptomycin, N2 growth supplement, epidermal growth factor (EGF) and fibroblast growth factors (FGF) (ThermoFisher Scientific, Waltham, MA, USA). U251_R132H and U251_IDH were cultured with DMEM with the supplement of 10%FBS.

### Metabolite Extraction

Cell pellets were collected at 48 hrs post-treatments and sonication by Misonix XL-2000 Ultra-liquid processor (Misonix Inc., Farmingdale, NY, USA) at 40 amps for 30 s. Following sonication, metabolite extraction was normalized and prepared using Methanol:Chloroform:water protocol and stored at –80 °C as previously described^16^.

### Cell Proliferation Assay (CCK-8)

15000 cells per well (Ts603 and GSC923) or 2000 cells (U251_R132H and U251_WT) per well were seeded at the round bottom 96-well plate for six repeats. After the treatment of drugs, the cells were continued cultured in a cell CO_2_ incubator at 37 °C. CCK8 reagent (10ul/well) was added into the cells at 72 hrs post-treatment and the absorbance at 450 nm was examined using a microplate reader.

### TBARS assay

0.5 × 10^6^ cells (Ts603 and GSC923) or 0.1 × 10^6^ cells (U251_R132H and U251_WT) per well were repeatedly treated with MF-438 for 3 days. The cells were collected at indicated time and lysis with cell lysis buffer 3 (Cat No: 895366). After acid treatment, the assay was followed instructions (R&D, Cat No: KGE013). The results were examined and read at 450 nm using a microplate reader. The protein concentration of each sample was used to normalize the data.

### Colony formation assay

0.5×10^5^ Ts603 or GSC923 were cultured in 0.5% low-melt temperature agarose gel. The cells were treated with 1uM MF-438 or DMSO or 4.3 μM DMT1 blocker 2 every 2∼3 days. The number and diameter of the colony were counted using the Celigo machine at day 0 and day 21.

### Flow cytometry analysis for iron metabolism

FerroOrange (Dojindo Inc, F374) was prepared by adding 35 µL of DMSO to a tube containing 24 µg of FerroOrange and dissolve the solution via pipetting to prepare a 1 mmol/L FerroOrange solution. Then, dilute the 1mmol/L FerroOrange solution with HBSS to prepare the 1 µmol/l FerroOrange working solution. Ammonium iron(II) sulfate 100mM dilute to 10mM by DDW. 10 mmol/L Ammonium iron (II) sulfate (10 uL) was used to stimulate the cells (The final concentration: 100 umol/L). The cells were incubated for 30 min in a 37°C incubator equilibrated with 95% air and 5% CO2. The cells were collected and washed with HBSS (1 mL) for three times. Then HBSS (1 mL) was added to the microcentrifuge tube and suspended by pipetting. 1 umol/L FerroOrange in HBSS (400 uL) was added to the cells and the cells were incubated for 15 min in a 37°C incubator equilibrated with 95% air and 5% CO2. Afte 15 min, the stained cells were passed through a cell strainer and analyze samples using a flow cytometer.

### Iron ELISA assay

After 3 days of treatment with DMSO (0.08%), TMZ (6.25uM), MF-438(1uM or 10nM), TMZ(6.25uM) + MF438(10nM), the same number of cells in each condition were stimulated with PBS or 100uM FeCl2 or 100uM FeCl3, respectively. Then, the cells were incubated at 37T for 30 min and vortex for every 5min. The cells were spin down at 300g for 5min and wash with PBS for 2 times. 50 mM NaOH was used for lysis the cells (400ul for 5×10^5 cells). 200ul lysates mixed with 200ul 10mM HCL and 200 uL iron releasing reagent (a mixture of 1.4 M HCl and 4.5% KMnO4). The reaction was run in 60T for 2 hrs. Iron Detection Reagent 45ul (6.5 mM Ferrozine (HY-137805,MCE), 6.5 mM neocuproine (HY-W004563, MCE), 2.5 M ammonium acetate (A16343.30, Thermo Scientific), and 1M ascorbic acid (A4544, Sigma) was added into the mixture at indicated time. The value was measured at OD550 after 30min incubation.

### Iron pool staining

Calcein AM staining buffer was prepared by adding 20 μL of the 1 mM calcein AM stock solution to 10ml HBSS. The cells were washed with HBSS for twice and stained using 500ul staining buffer at 37T for 30min. The supernatant was aspirated, and the cells were fixed with 2% PFA at RT for 15min. After 2 washed with PBS, Nuclei staining with Hoechst 3342 at RT for 5min. The slides were prepared by mounted and sealed.

### Western Blot

The cells were collected and lysis using SDS lysis buffer. Equal amounts of protein were loaded in each lane for 4-15% Bio-Rad Mini Protean TGX gel and then transferred to polyvinylidene difluoride membranes. The membranes were washed with blotting buffer (1× PBS containing 0.1% Tween20) and then blocked for 60 min in blotting buffer containing 10% low-fat powdered milk. Membranes were washed 3 times with blotting buffer, incubated at 4° C overnight with primary antibody (1:1000) containing 5% lowfat powdered milk, and incubated with secondary antibody (1:1000) at room temperature for 60 min. Target proteins were developed with Bio-Rad Clarity Western ECL substrate. GPX4 (Cat No. 52455), DMT I (Cat No. 15083), SCD1(Cat No. 52455S), FTHI (Cat No. 4393S), beta-actin (Cat No. 4970), was purchased from cell signaling tech Inc. SCD1 (Cat No. ab19862) and Ferittin Light(Cat No. ab69090) were purchased from Abcam. The blots were detected with BioRad ChemiDoc Touch imaging system. The relative expression of proteins was normalized to beta-actin and analyzed using Image J.

### TMZ and MF-438 Synergy Assay

Cells were seeded at a density of 15,000 cells per well (D0) in 96-well plates and allowed to attach overnight. TMZ was serial diluted at 3.12uM, 6.25uM, 12.5uM, 25uM, 50uM, 100 μM. MF-438 was prepared by serial dilution at 1nM, 10nM, 25nM, 50nM, 100nM, 250nM, 500nM, 1uM. to generate a dose–response matrix. Cells were treated with the combination of MF-438 and TMZ or vehicle control to generate a dose–response matrix. Then the cells were incubated at 37 °C with 5% CO₂ for 72 h. Cell viability was measured using cck8 assay according to the manufacturer’s instructions. The results were analyzed by Synergy Finder R-package.

### Animal Experiments

Six-week-old female SCID (NOD, Charles River) mice were i.c., injected with 1.8

× 10^5^ patient-derived glioma stem cells TS603 cells (day 0). The mice were randomly allocated (8∼10 mice/group) for 3 groups. Then the mice were gavaged with 10% DMSO as control, TMZ (50 mg/kg), MF-438 (1.25 mg/kg) + TMZ (50mg/kg) on day 15, followed by a second treatment on day 30. The endpoint of the study was determined by monitoring the body condition of the mice. Tumor presence was examined and confirmed by histopathological analysis of brain tissues. Kaplan–Meier analysis was performed to evaluate disease outcomes. The animal protocol NOB-008 was approved by the National Cancer Institute (NCI) Animal Use and Care Committee.

### Knockdown of SCD1 and SFA rescue experiments

2000 cells (U251_R132H and U251_WT) per well were seeded at the round bottom 96-well plate for six repeats. The cells were transfected with a pool of 3 SCD1-target specific siRNAs (OriGene, SR321692) or siRNA control (OriGene, SR30004) using Lipofectamine RNAiMAX transfection reagent (ThermoFiher, 13778150). At two days post-transfection, the cells were treated with vehicle control, or SFA 10ug/ml. Cell viability was determined by cck8 assay.

### SFA induced synergy with MF438

TS603 cells were treated with 1uM MF438 or DMSO with or without 10uM BSA or 10uM SFA and the relative cell number was measured after 24 hours. U251 WT or U251R132H cells were also treated with 1uM MF438 and 10 uM SFA (C18:0, C16:0, C18:1, C16:1) were added.

### RNA Sequencing Experiment and Data Analysis

Total RNA was extracted using RNeasy spin columns (Qiagen, Cat No:74136) with RNase-free DNase (Qiagen, Chatsworth, CA) from treated glioma cells. RNA quality was assessed by a Nano Drop 2000 Bioanalyzer, and RNA-Seq was performed by NIH Frederick NGS core facility. RNA-Seq analysis was conducted using R packages: clusterProfiler, enrichplot, and Deseq2. Feature lists were filtered by inter-quartile range (bottom 5 %) and quality control precision (<25 % RSD) prior to sum-normalization and auto-scaling. Inclusion in lists of significantly dysregulated metabolites was based on volcano plot results (p < 0.05; FC > 2.0). Heat maps were generated from sum-normalized (but not auto-scaled) data for relevant features.

### Co-IP between DMT1 and SCD

Co-IP assays were performed using antibody conjugated Dynabeads™ magnetic beads following the manufacturer’s instructions (ThermoFisher, 14321D). Proteins were eluted from magnetic beads in SDS buffer supplemented with SB buffer by shaking at 100 °C for 5 min.

### NADP^+^/NADPH Ratio Measurements

2.0 × 10⁶ cells were lysed in NADP/NADPH extraction buffer and filtered through a 10 kDa MWCO filtration tube. To distinguish total NADP⁺/NADPH from NADPH, samples were either heated at 60 °C for 60 min or left unheated, respectively. Samples were then transferred to a 96-well microplate and incubated with the working solution at 37 °C for 60 min. NADP⁺/NADPH and NADPH levels were quantified following the manufacturer’s instructions (DoJinDo, N510-10).

## RESULTS

### D-2HG–induced SCD1 expression creates a metabolic vulnerability in IDH1^MUT^ glioma

SCD1 is the rate limiting step of lipid desaturation and a recognized metabolic dependency in multiple cancers. While SCD1 inhibition has shown anti-tumor efficacy in diverse contexts, the potential of SCD1 targeting in glioma has remained poorly defined. To assess the potential contribution of SCD1 to IDH1-mutant gliomagenesis, we analyzed its expression in both publicly available and patient-derived datasets. SCD1 expression was significantly enriched in IDH1-mutant oligodendroglioma compared with IDH1-wild-type glioblastoma (Fig. 1a) in the TCGA cohort and was similarly elevated in IDH1-mutant patient samples (Fig. 1b). Importantly, overexpression of IDH1^MUT^ alone was sufficient to elevate SCD1 protein levels in both malignant (U251 glioma) and normal human astrocyte models (Fig. 1c), indicating that SCD1 upregulation is a direct, cell-autonomous consequence of mutant IDH1 activity rather than a secondary adaptation to transformation. Unbiased transcriptomic profiling revealed that D-2HG elicits one of the most robust transcriptional responses in the lipid metabolic network, with SCD1 emerging as a prominently upregulated transcript in IDH1^WT^ cells exposed to D-2HG (Fig. 1d). This induction was not limited to transcript levels; D-2HG treatment produced a concordant and substantial increase in SCD1 protein expression, demonstrating a tightly coupled transcriptional and translational response (Fig. 1e).

**Figure 1.**
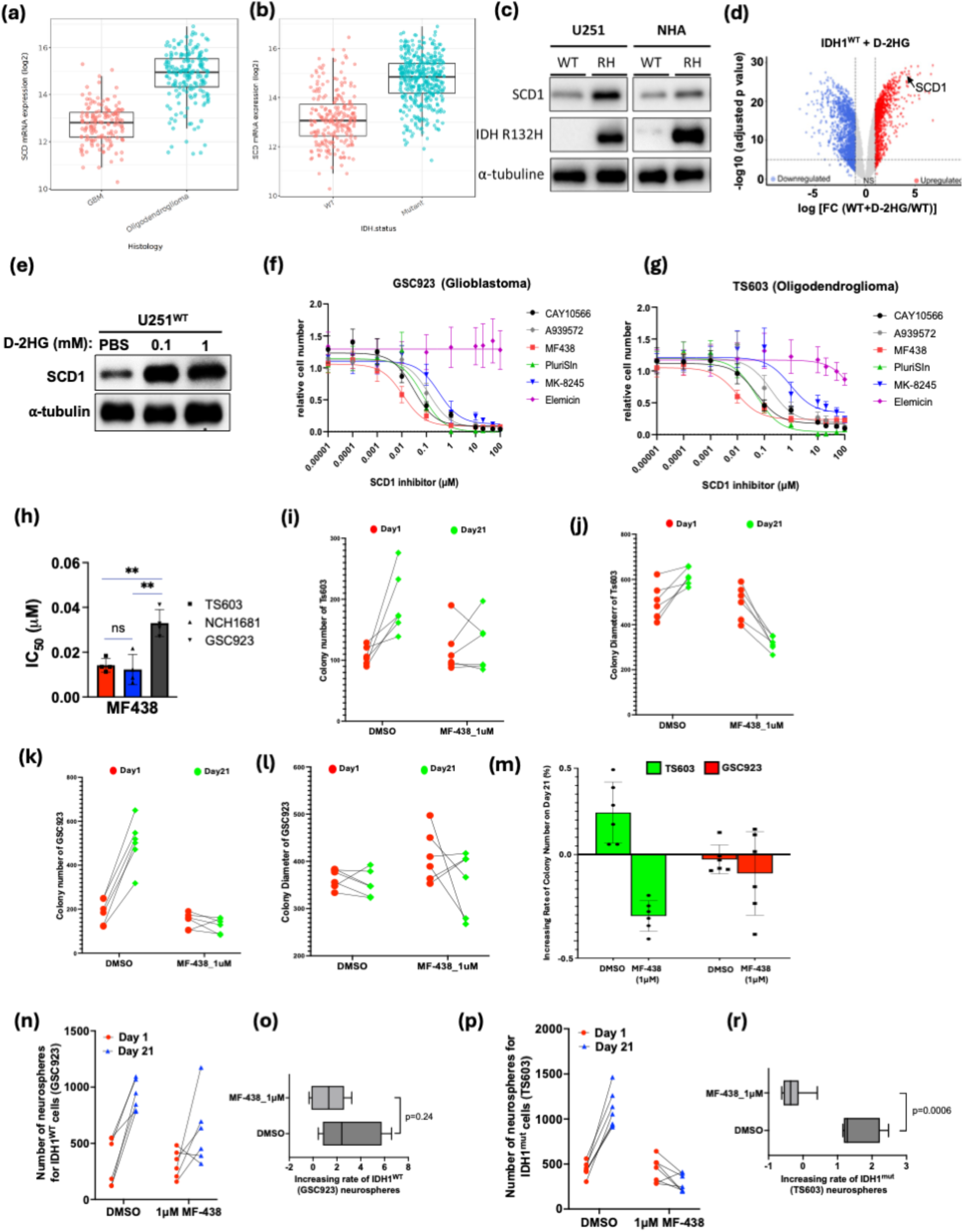
SCD1 expression is regulated by D-2HG, and its pharmacological inhibition leads to significantly reduced cell growth and colony formation. a) and b). mRNA levels of SCD1 from TCGA show an increased expression of this transcript in oligodendroglioma when compared with GBM (a) and in IDH1^MUT^ when compared with wild type (b). c). SCD1 protein levels increase upon IDH1^mut^ overexpression in U251 cells and normal human astrocytes. d) D-2HG increases expression of SCD1 (volcano plot) when added to IDH1^WT^ U251 cells. e) D-2HG addition to U251 IDH1^WT^ cells increase the expression of SCD1 at the protein levels as well. f) and g CCK-8 assay for multiple inhibitors of SCD1 tested in IDH1^WT^ patient derived cell line, GSC923 and in IDH^MUT^ patient derived cell line, TS603. Each curve is the result of three independent replicates. The data is fitted using drug response preset function from Graph Prism. h) IC_50_ value extracted from the CCK-8 is plotted for the compound MF-438 which was the most potent, for three cell lines, TS603 (IDH1^mut^ oligodendroglioma, GSC923 (IDH1^WT^ GBM), and NCH1681 (IDH1^mut^, astrocytoma). i-t) Colony forming assays MF 438 affects the rate of colony formation in TS603 by affecting the colony number (i) and its diameter (j) measured at day 1 and day 21. Colony forming assays MF 438 does not affect the rate of colony formation in GSC923 by measuring at the colony number (k) or their diameter (l) at day 1 and day 21. The rate of neurosphere formation was significantly altered in TS603 but was not significant in GSC923.

Having confirmed D-2HG-induced upregulation of SCD1, we next investigated whether pharmacological blockade of SCD1 could suppress glioma growth. Treatment with nine structurally distinct SCD1 inhibitors revealed a striking, genotype-specific effect: IDH1^MUT^ glioma cells were profoundly sensitive to SCD1 inhibition, whereas IDH1^WT^ GBM cells exhibited lower sensitivity via higher IC_50_ (Fig. 1f-1h). Preferably, SCD inhibitors did not affect normal human astrocyte cell growth, strengthening the safety of SCD inhibitor treatment for glioma. (Fig. 1d and 1e, Supplemental Fig. S1a-S1c). The IC_50_ was significantly lower in IDH1^MUT^ cell lines (Fig. 1h). Additionally, IDH1^MUT^ gliomas exhibited significantly greater sensitivity to multiple SCD1 inhibitors than IDH1^WT^ GBMs, as demonstrated by a reduced number of colonies in clonogenic assays conducted in soft agar for both patient-derived cell lines as well as isogenic glioblastoma cell lines overexpressing IDH1 mutation (Fig. 1i-1j). Interestingly, the rate of colony formation was not affected by SCD1 inhibition for the patient-derived IDH1^WT^ cells but significantly decreased for IDH1^MUT^cells (Fig.1m-1r).

Additionally genetic depletion of SCD1 using siRNA significantly reduced proliferation, with IDH1^MUT^ cells exhibiting a greater sensitivity to SCD1 loss than IDH1^WT^ cells, suggesting an increased dependence on SCD1-mediated lipid desaturation in the IDH1^MUT^ background (Supplemental Fig. S2d-S2e).

To determine whether the growth inhibition was linked to altered fatty acid composition, cells with SCD1 knockdown were supplemented with either saturated fatty acids (SFAs) or monounsaturated fatty acids (MUFAs). Addition of the SFAs palmitate (C16:0) and stearate (C18:0) further decreased cell viability in both IDH1^WT^ and IDH1^MUT^ cells, indicating that excess saturated fatty acids exacerbate the effects of SCD1 loss. In contrast, supplementation with the MUFAs palmitoleate (C16:1) and oleate (C18:1) restored relative cell numbers, demonstrating that the growth defect induced by SCD1 depletion is largely attributable to a deficiency in monounsaturated fatty acids. (Supplemental Fig. S2f-S2g).

To further validate this mechanism, cells were pretreated with the SCD1 inhibitor and subsequently supplemented with increasing concentrations of MUFAs. Dose-dependent rescue of cell viability was observed in both IDH1^WT^ and IDH1^MUT^ cells, confirming that exogenous MUFAs can compensate for the loss of endogenous SCD1 activity (Supplemental Fig. S2h-S2k). Representative neurosphere images corroborated these findings, showing impaired sphere growth following SCD1 inhibition and restoration of neurosphere formation upon MUFA supplementation (Supplemental Fig. S2l).

Western blot analysis revealed that treatment with the SCD1 inhibitor increased SCD1 protein expression, suggesting activation of a compensatory feedback response to reduced desaturase activity. In contrast, supplementation with MUFAs decreased SCD1 expression, consistent with feedback inhibition by the end products of SCD1-mediated fatty acid desaturation (Supplemental Fig. S2m). Together, these findings establish that glioma cells depend on SCD1-derived MUFAs for proliferation and survival, with IDH1^MUT^ cells displaying heightened sensitivity to disruption of this pathway.

### Loss of buffering capacity of IDH1^MUT^ cells for saturated fatty acids explained their sensitivity to SCD1 inhibition

To investigate the mechanism of SCD1 inhibition, we used MF-438 for the rest of the study, the most potent inhibitor identified from the screen of nine commercially available compounds, which exhibited the strongest suppression of colony formation in IDH1^MUT^ cells while having no significant effect on the rate of colony forming of IDH1^WT^ cells (Fig 1). To test whether impaired handling of saturated fatty acids in IDH1^MUT^ cells is linked to their dependence on lipid desaturation, we evaluated the impact of SCD1 inhibition on saturated lipid accumulation, mRNA, and cell viability. Targeted lipidomic assay show no significant difference in the substrates of the products of SCD1 in IDH1^WT^ cell line (GSC923) while a significant drop of monounsaturated fatty acids oleic (C18:1) and palmitoleic (C16:1) were observed in IDH1^MUT^ cell line (TS603), confirming a mutant-specific dependence on SCD1-mediated desaturation (Fig 2a-2d). These findings confirm on-target inhibition of SCD1 and demonstrate remodeling of cellular lipid composition following treatment. Interestingly, MF-438 treatment elicited widespread remodeling of phospholipid composition in IDH1^WT^ cells (GS923), with strong accumulation of saturated fatty acids, DG, PE, PC, ceramide, SM, and PS species. These lipidomic changes were less pronounced or missing all together in IDH1^MUT^ (TS603) cells, suggesting the ability of IDH1^WT^ cell line to buffer these lipids by placing them in downstream metabolites (Fig. 2e). To determine the functional importance of SCD1, we assessed the effects of acute SCD1 depletion on cell viability and evaluated whether exogenous fatty acid supplementation could rescue the resulting phenotype.

**Figure 2.**
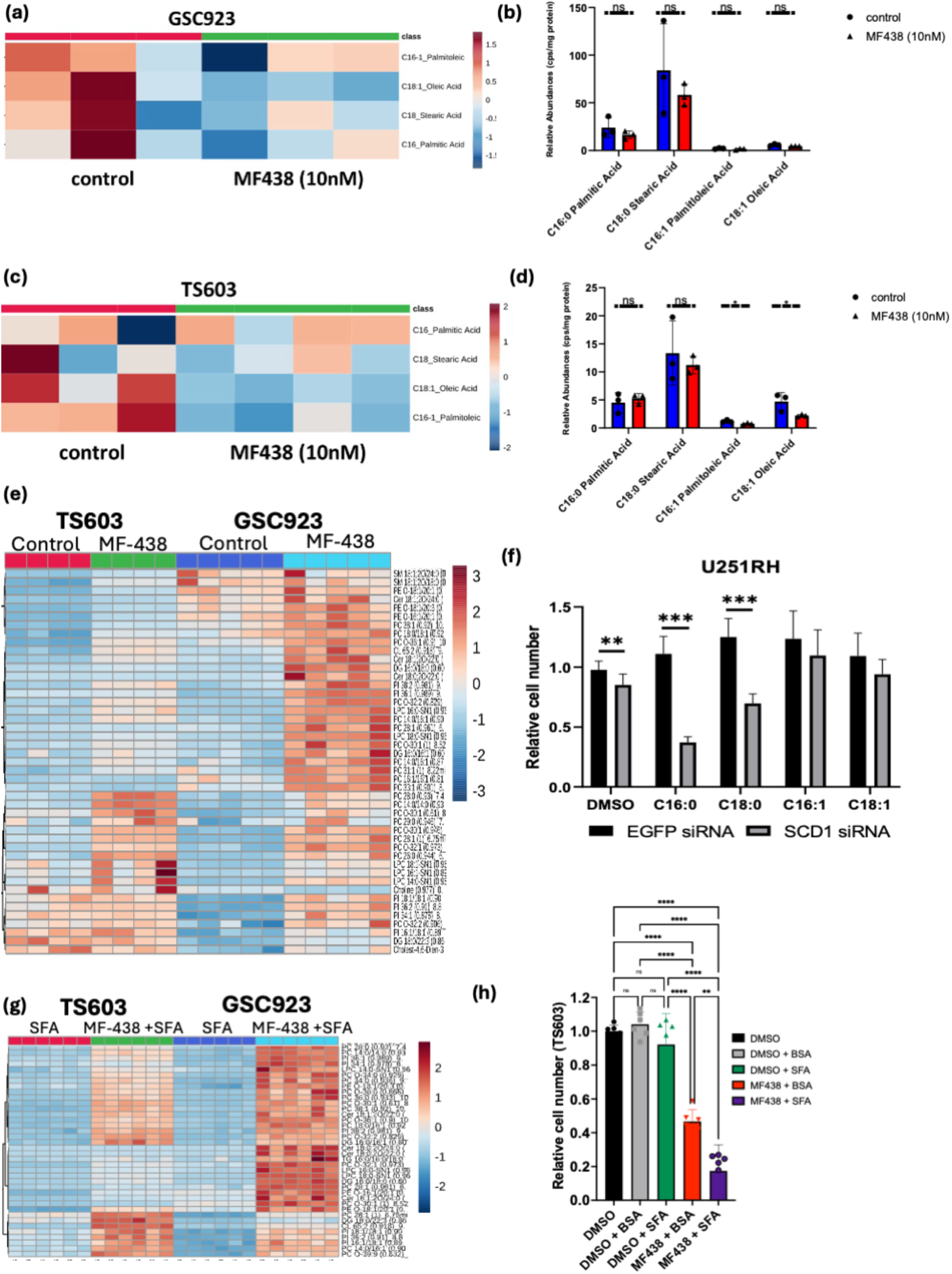
SCD1 inhibition induces saturated fatty acid accumulation, lipidomic remodeling, and decreased cell proliferation. a)-d) Heatmap and corresponding bar graph of SCD1 inhibition by MF-438 in IDH1^WT^ cells (GSC923) a) and b) and IDH1^MUT^ cells (TS603), c) and d). e). Heatmap of downstream PCs, PEs, LPCs, Ceramides, DGs, PIs, sphingomyelins (SM) enriched in SFAs highly upregulated in GSC923 and not so much in TS603. f).Bar graphs of siRNA of SCD1 showing the effects on proliferation and the further effects of SFA addition of MUFA addition. g). Heatmap of lipid affected by MF-438 and SFA addition in TS603 and GSC923 showing buffering capacity of GSC923. h). The cumulative effect of SFA and MF-438 on proliferation of TS603.

siRNA-mediated knockdown of SCD1 significantly impaired cell viability. This effect was reversed by supplementation with palmitic acid (C16:0) or stearic acid (C18:0), suggesting that SCD1-regulated lipid metabolism plays a critical role in sustaining proliferative capacity (Fig. 2f). These data align with our next observation that exogenous SFA supplementation exacerbates the growth-inhibitory effects of MF-438 in IDH1^MUT^ cells. To validate that the loss of buffering capacity of IDH1^MUT^ cells is correlated with increased sensitivity of these cells to SCD1 inhibitor, we pretreated the cells (IDH1^MUT^ and IDH1^WT^) with SCD1 inhibitor and added on top of that either monounsaturated fatty acids or saturated fatty acids.

Monounsaturated fatty acid supplementation partially rescued the MF-438–induced phenotype, demonstrating that the vulnerability reflects impaired desaturation capacity rather than global lipid depletion (Supplemental Fig. 2h-2m). The accumulation of SFAs following SCD1 inhibition likely exceeds the cellular capacity for lipid buffering and storage, leading to lipid metabolic stress (Fig. 2h). Conversely, IDH1^WT^ cells appear to retain a greater ability to sequester or metabolize excess SFAs into downstream lipid pools, thereby mitigating the deleterious effects of SFA accumulation (Fig 2g).

Furthermore, RNA seq analysis show lipid that droplet organization genes (SQLE, CHKA, AUP1) were coordinately upregulated in TS603 after treatment with MF-438 together with increased lipid processing genes and lipid transport genes (Supplemental Fig. 3).Together, these data indicate that IDH1^MUT^ cells rely on SCD1 activity to buffer excess saturated fatty acids, and that loss of this buffering mechanism contributes to their heightened sensitivity to SCD1 inhibition.

### Iron homeostasis disruption and labile iron pool alterations as mechanistic drivers of MF-438-induced ferroptosis

Since SCD1 is an Fe^2+^ dependent desaturase, we looked at the correlation between SCD expression and iron import protein SLC11A2 (DMT1) in TCGA specifically for IDH1^MUT^ oligodendroglioma patients. A positive correlation as measure by Pearson’s product-moment correlation was observed in this subset of glioma suggesting a link between the import of iron and SCD1 (Fig. 3a). The Pearson correlation coefficient was lower in IDH1^WT^ GBM suggesting a less pronounce effect than in IDH1^MUT^ oligodendroglioma. Having this positive association between DMT1 and SCD1, we next measured the labile iron pool using confocal microcopy in DMSO and MF-438 treated TS603 cells (Fig. 3b-3c). MF-438 decreased the labile iron pool in IDH1^MUT^ cells, and this was not further affected by the addition of SFAs. Consistent with the lower association data between DMT1 and SCD1 in GBM, addition of MF-438 did not alter the labile iron pool for the IDH1^WT^ cells. Next, we measured ferrous iron (Fe²⁺) content using FerroOrange and flow cytometry which showed a decrease in the count of cell containing Fe^2+^ of upon the addition of MF-438 in IDH1^MUT^ cells (Fig. 3d). Since iron levels are connected to ROS and ferroptosis we next measured NADP+/NADPH ratios as a proxy of ROS levels based on the well-known assumption that (NADP+/NADPH) redox couple serves as a substrate or cofactor for many enzymes to maintain cellular redox homeostasis. Interestingly, we measured a decreased in this ratio as a function of MF-438 addition, which would suggest increase reductive stress, and this was in contrast with the increase in TBARS which is an indirect measure of ROS for IDH1^MUT^ cells. Contrary, IDH1^WT^ cells had a similar increase in reductive stress but a decrease in ROS (Fig. 3e-3f). At the protein levels, MF-438 decreased DMT1 levels, GPX4 and FTH1, which agrees with a decreased iron import and transport in MF-438 together with a decrease anti-ferroptosis response (Fig. 3g). This data also matches the marked decrease in cell viability in TS603 in presence of iron chelator molecule, deferasirox (Fig. 3h-3i). Overall, these trends were recapitulated at the RNA levels in MF-438 treated cells (Fig. 3j). Together these data highlight a link between MF-438 treatment and iron response inhibited, cells might shift toward a “low iron utilization” phenotype, turning down iron import (DMT1) and ferritin (FTH1) as a protective adaptation.

**Figure 3.**
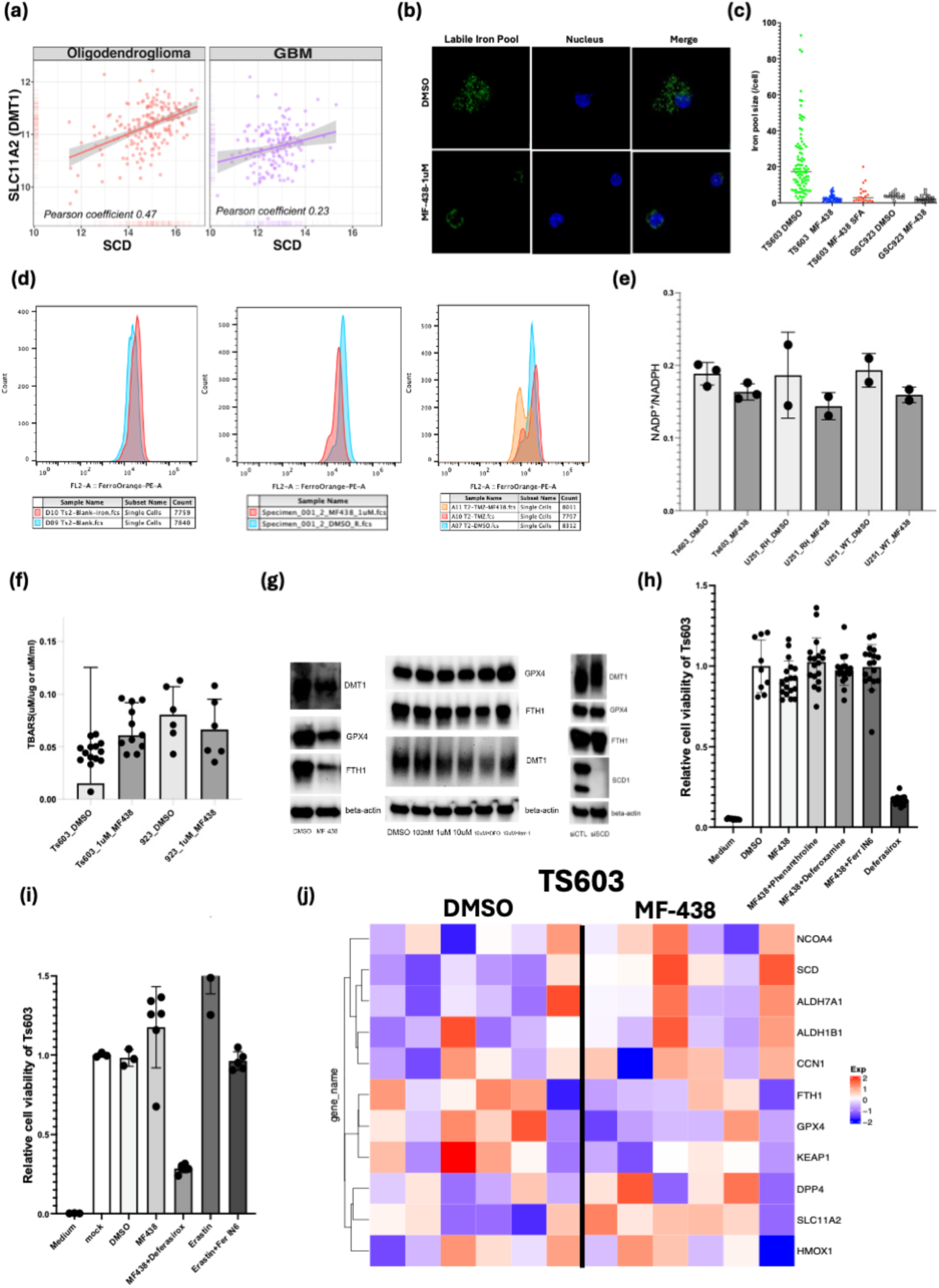
Iron homeostasis disruption and labile iron pool expansion as mechanistic drivers of MF-438-induced ferroptosis. a) Pearson correlation analysis of mRNA for SLC11A2 (DMT1) levels and SCD1 levels in oligodendroglioma (left) versus glioblastoma (right). c) and d) Confocal microscopy of labile iron pool (green) and nuclei (blue) in absence and presence of MF-438 and its quantification. e) Flow cytometry analysis of Fe^2+^ levels in absence and presence of MF-438. e) NADP+/NADPH ratio of three cell lines in presence of MF-438. f) TBARS measurements in presence MF-438. g) Western Blot analysis of DMT1, GPX4 and FTH1 in presence of MF-438. h) and i). Cell viability of TS603 in presence of addition, neutral lipid metabolism genes (LPIN1, LDLR, PNPLA3, ACSL1), intracellular lipid transport genes (TMEM41B, OSBP, STARD4), and lipid droplet organization genes (SQLE, CHKA, AUP1) were coordinately upregulated in TS603 after treatment with MF-438 (Supplemental Fig. 3).

### Blocking DMT1-dependent iron uptake mimics MF-438 efficacy in IDH1-mutant glioma cells

Given that increased iron uptake via DMT1 emerged as a prominent signature of MF-438 treatment, we next investigated the role of iron homeostasis in modulating cell viability. Supplementation with Fe²⁺ at concentrations up to 100 μM did not affect the viability of either IDH1-mutant or IDH1-wild-type cells (Fig. 4a–b). However, Fe²⁺ treatment significantly increased DMT1 protein expression (Fig. 4c). Notably, pharmacologic inhibition of DMT1 markedly reduced the viability of IDH1^MUT^ (TS603) cells, as demonstrated using two independent DMT1 inhibitors and escalating inhibitor concentrations (Fig. 4d–f). Co-immunoprecipitation studies further revealed an interaction between DMT1 and SCD1 (Fig. 4g). Consistent with these findings, DMT1 inhibition significantly decreased both colony formation and neurosphere growth in IDH1-mutant cells (Fig. 4h–k), whereas these effects were not observed in IDH1-wild-type cells (Fig. 4l–m). These findings suggest that IDH1^MUT^ glioma cells exhibit a heightened dependence on DMT1-mediated iron uptake.

**Figure 4.**
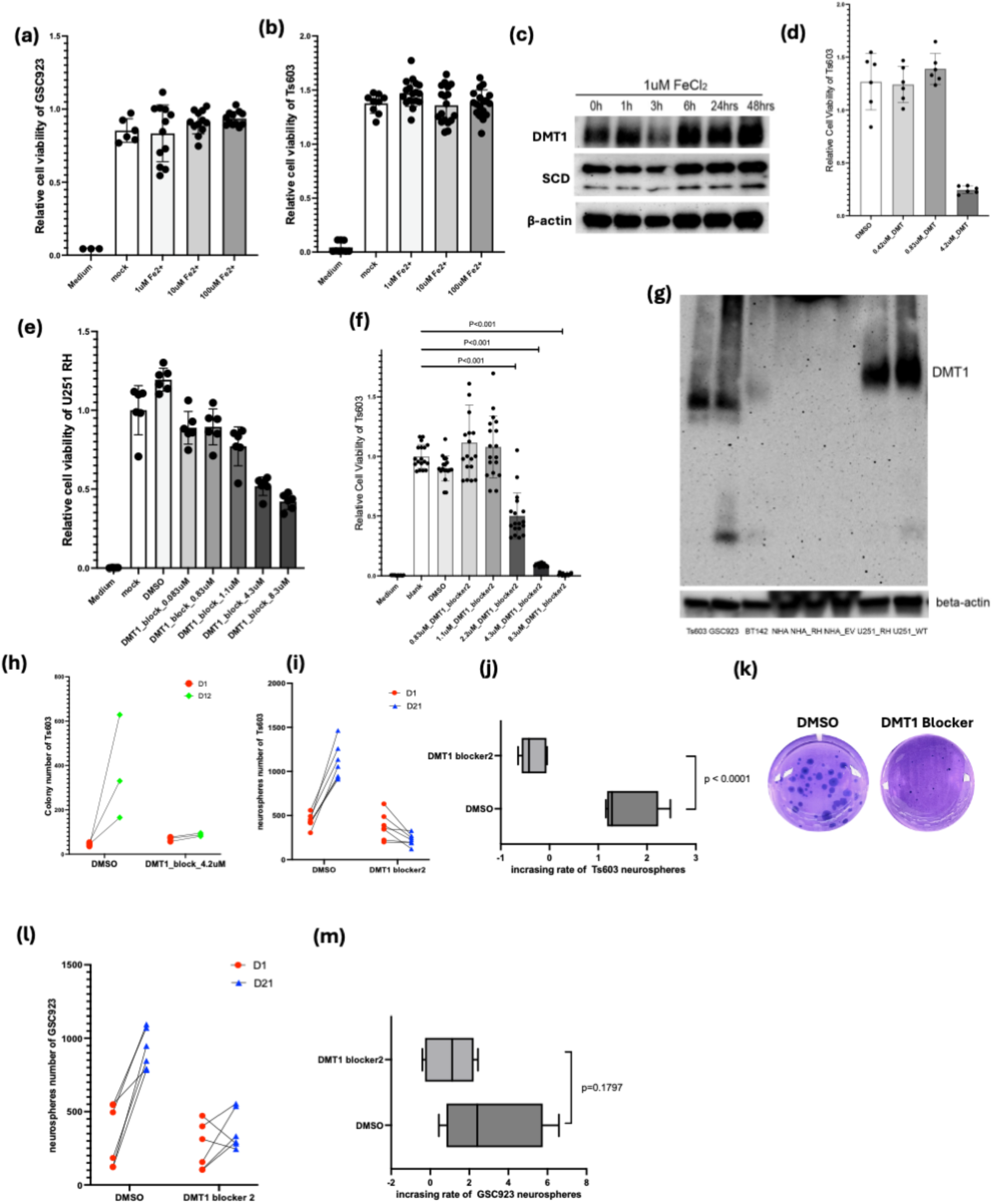
DMT1-blocking mimics the effects of MF-438. a) and b) Cell viability of GSC923 and TS603 respectively, as a function of increasing Fe^2+^ concentrations. c) Western Blot of DMT1 and SCD1 proteins as a function of increasing Fe^2+^ concentrations d)-f). TS603 cell viability as a function of DMT1 blocker 1 and blocker 2. g) Co-IP of SCD1 and DMT1 in different cell lines. h)-k). Colony forming assay of TS603 in presence of DMT1 blockers. l)-m). Colony forming assay of GSC923 in presence of DMT1 blocker 2.

### SCD1 inhibition markedly sensitized glioma cells to temozolomide (TMZ) in vitro by activating both necroptosis and ferroptosis

Since temozolomide (TMZ) is one of the standards of care for the oligodendroglioma that might be sensitized to the increased oxidative stress, we wanted to combine the MF-438 treatment with TMZ and see if we can enhance its efficacy. Combined MF-438 and TMZ resulted in a pronounced loss of viability and clonogenic potential Fig. 5a-5c. Like SCD1 inhibition, this effect was more pronounced in IDH1^MUT^ glioma when compared with IDH1^WT^, where the combination synergistically suppressed TS603 neurosphere growth (p=0.0326) We then perform RNAseq analysis in both IDH1^MUT^ and IDH1^WT^ in the presence of MF-438 and MF-438 + TMZ (combo) to identify which mechanisms of cell death are the most predominant. Interestingly, MF438+TMZ activated the intracellular lipid transport pathways and neutral lipid metabolic processes (Fig. 5g and Supplemental Fig. 3). Furthermore, the combination treatment also triggered an integrated ER stress response (ATF3, DDIT3, IRF1, CDKN1A, GADD45B) and increased the mRNA for proteins involved in ferroptosis and necroptosis (Fig. 5h-5i). Together, these findings show that SCD1 inhibition disrupts lipid homeostasis and iron handling in a manner that compromises redox buffering and DNA repair capacity, thereby lowering the threshold for TMZ-induced cytotoxicity and leading to ferroptosis and necroptosis cell death.

**Figure 5.**
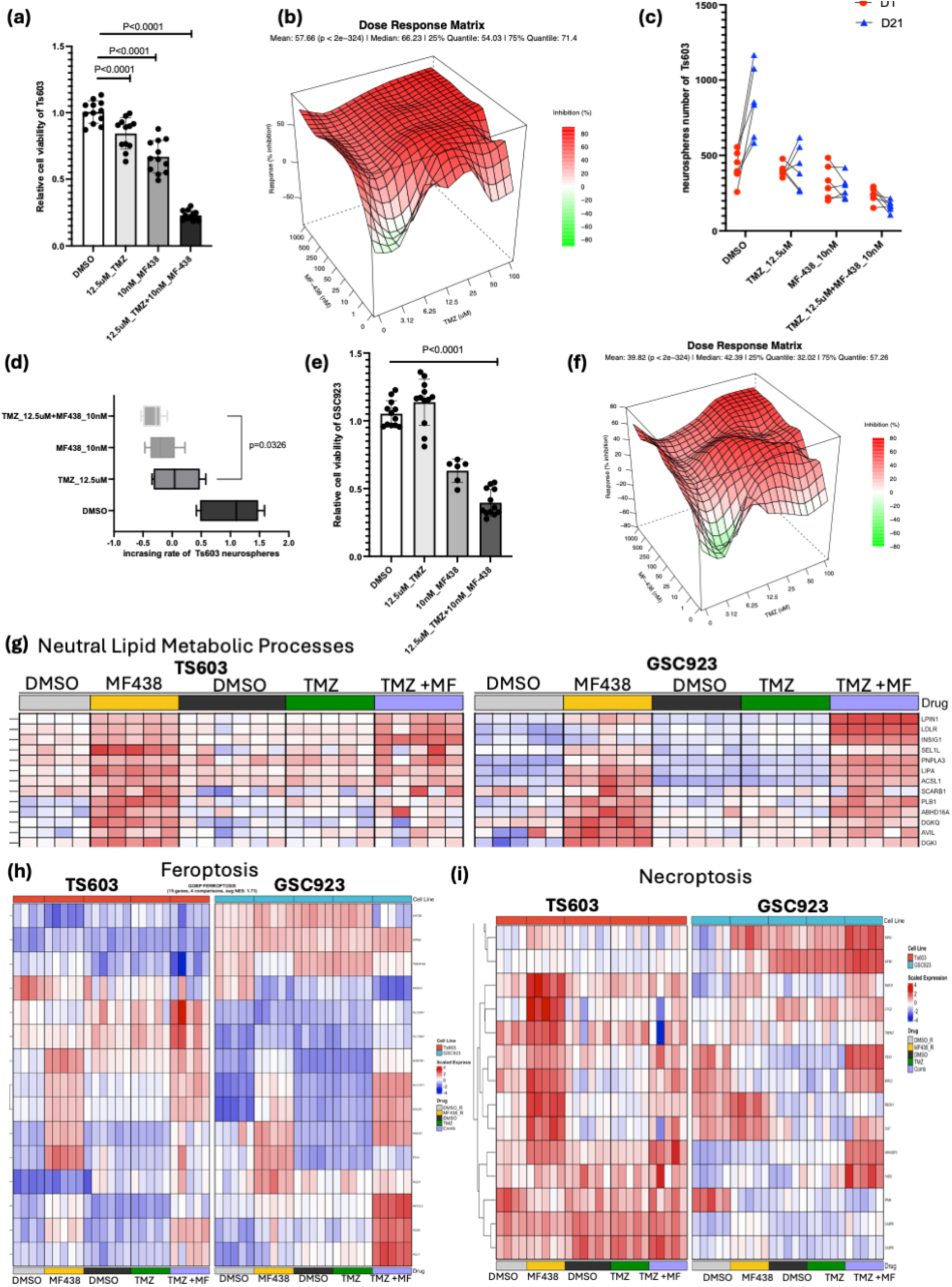
Synergetic effects of MF-438 and TMZ in vitro. a)-f) MF-438+TMZ combination demonstrates superior in vitro cytotoxicity, neurosphere suppression, and synergistic dose-response more pronounced in IDH1^MUT^ cell line. g) The combination also affects the neutral lipid metabolic processes, increases ferroptosis (h) and necroptosis markers (i).

### MF-438+TMZ combination therapy extends median survival and reduces Ki-67 proliferation in IDH1-mutant TS603 glioma mouse model

Finally, we tested whether the combination therapy is more efficient in vivo in an intracranial model of IDH1-mutant oligodendroglioma. Tumors were implanted intracranially in SCID mice. At day 15, the treatment for all the groups was started in 2 dosing schedules to match the clinical dosing. In this context, we designed the groups to match the TMZ dosing schedule, which means that we only had 10 doses of SCD1 inhibitor together with 10 doses of TMZ, and in the single-treatment arm we matched this dosing schedule (Fig 6a).

**Figure 6.**
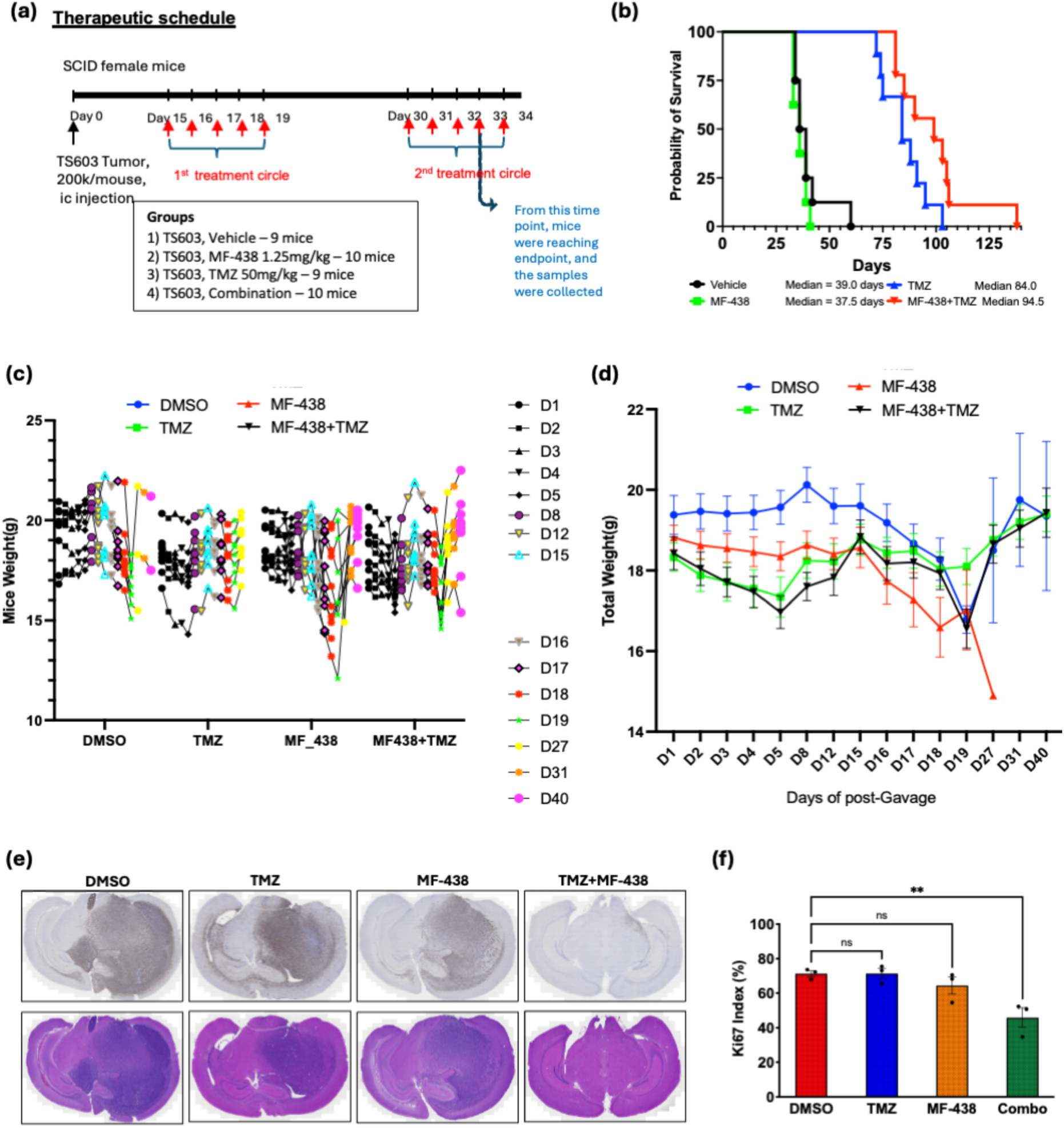
Synergetic effects of MF-438 and TMZ in vivo. a) Therapeutic schedule of mice harboring TS603 cells intracranially. b) Survival of mice treated with only 10 doses of MF-438, TMZ and TMZ+ MF-438. c)Mice weight during the experiment for all the groups. d) Total weight as a function of days for all the groups. e) Immunohistochemistry for Ki67 and the corresponding H&E stain. f) Quantification of Ki67 positive cells.

As expected, the SCD1 inhibitor alone did not improve survival because we did not dose the mice continuously; we only gave 10 doses. The largest survival benefit was observed with TMZ, as expected and matching the clinical findings in oligodendrogliomas (Fig. 6b). Interestingly, the combination therapy had a significant survival benefit over TMZ alone, which was also reflected by a decrease in the proliferation marker Ki67 (Fig. 6e-6f).

These data demonstrate that combining an SCD1 inhibitor with temozolomide (TMZ) provides a greater therapeutic benefit than TMZ alone in an intracranial IDH1-mutant oligodendroglioma model. While short-term SCD1 inhibition by itself did not improve survival, the combination treatment significantly extended survival and reduced tumor cell proliferation, as indicated by lower Ki67 levels. These findings suggest that SCD1 inhibition can enhance the antitumor efficacy of TMZ and may represent a promising strategy for improving treatment outcomes in IDH1-mutant oligodendroglioma.

## DISCUSSION

SCD1 is increasingly recognized as an important regulator of cancer biology, influencing processes such as cancer stemness and immune responses^21,31,34,35^. Despite its established role in other malignancies, its function in brain tumors remains insufficiently understood, highlighting the need for further investigation and supporting the rationale for exploring SCD1 as a potential therapeutic target in brain cancer. Herein, we identify that a subset of gliomas with an IDH1 mutation are more sensitive to SCD1-based pharmacological targeting and link this inhibition to iron homeostasis and two mechanisms of cell death, ferroptosis and necroptosis.

We found that treatment with MF-438 alone reduced monounsaturated fatty acid (MUFA) levels while increasing markers of lipid and endoplasmic reticulum (ER) stress. Mechanistically, SCD1 inhibition altered iron homeostasis, leading to reduced expression of the iron transporter DMT1 and the iron storage protein FTH1. These changes were accompanied by a decrease in intracellular iron and are consistent with impaired iron uptake and increased susceptibility to oxidative stress.

We also described a physical interaction between DMT1 and SCD1, which together with the selective sensitivity of IDH1-mutant cells to DMT1 inhibition, supports a functional link between iron metabolism and lipid desaturation. These findings suggest that iron availability plays a previously underappreciated role in regulating lipid homeostasis. Because desaturases such as SCD1 and FADS1/2 require iron as a cofactor, reduced DMT1 activity is expected to impair fatty acid desaturation, limiting the conversion of saturated fatty acids into MUFAs. The resulting loss of MUFAs and accumulation of saturated fatty acids necessitates metabolic adaptation. This differential metabolic response between IDH-mutant and IDH-wild-type cells was particularly striking. While IDH1^MUT^ cells depleted MUFAs from structural phospholipids such as PCs and PEs, IDH1^WT^ cells preferentially depleted MUFA-containing triacylglycerols. This pattern suggests fundamentally different strategies of MUFA utilization, with IDH1^MUT^ mutant cells drawing on membrane lipid pools and IDH1^WT^ cells relying more heavily on neutral lipid reserves. Unlike IDH1^WT^ cells, IDH1^MUT^ cells failed to mount an effective SFA lipid-buffering response. In contrast, direct SCD1 inhibition in IDH1^MUT^ cells reproduces the buildup of saturated fatty acids without inducing triglyceride storage, leaving phospholipid incorporation as the primary outlet. This selective rerouting likely increases membrane rigidity, and we postulate that might contribute to the heightened sensitivity of IDH1^MUT^ cells to MF-438. These findings align with our previous work demonstrating that IDH-mutant glioma cells preferentially channel monounsaturated fatty acids into ER and Golgi membranes, resulting in membrane abnormalities and ER dysfunction. Together, these results suggest that altered MUFA partitioning is a key feature of the metabolic rewiring associated with IDH1 mutations.

RNA sequencing analyses revealed that MF-438 led activation of ER stress pathways, as reflected by increased INSIG1 and SEL1L expression. In IDH1^MUT^ cells, MF-438 selectively reduced polyunsaturated phosphatidylcholine species and was associated with downregulation of GPX4, FTH1, and KEAP1, together with upregulation of NCOA4, SLC11A2, ALDH7A1, and DPP4. These transcriptional changes are indicative of ferroptosis priming through ferritinophagy-mediated remodeling of iron metabolism and agree with our iron pool measurement. In IDH1^WT^, MF-438 treatment led to concurrent induction of genes involved in neutral lipid metabolism (LIPN1, LDLR, PNPLA3, ACSL1), intracellular lipid trafficking (TMEM41B, OSBPL family members, and STARD4), and lipid droplet organization (SQLE, CHKA, and AUP1) suggests activation of compensatory lipid-buffering and transport pathways in response to SCD1 inhibition.

Unexpectedly, our findings reveal that DMT1 plays a critical role in supporting SCD1 function, suggesting a functional interplay between iron metabolism and lipid desaturation. These results further indicate that modulation of intracellular iron levels may represent a mechanism for regulating SCD1 activity, highlighting iron availability as a potential determinant of lipid metabolic homeostasis. We propose that SCD1 inhibition promotes a “low iron utilization” state characterized by reduced iron import and storage, representing a protective adaptation that links iron metabolism to lipid homeostasis. Collectively, these observations indicate that iron uptake, through its role in sustaining desaturase activity, is a critical determinant of fatty acid partitioning between structural and storage lipid pools and thereby shapes cellular responses to metabolic stress.

Based on our findings that SCD1 inhibition lowered the labile iron pool and decreased expression of FTH1 and DMT1, changes consistent with reduced iron uptake and heightened susceptibility to oxidative stress, we combined SCD1 inhibition with temozolomide (TMZ), the current standard of care. Our findings demonstrate that inhibition of SCD1 significantly enhances the antitumor activity of TMZ in glioma cells, particularly in the context of IDH1 mutations. While TMZ remains a cornerstone of therapy for oligodendroglioma, resistance to treatment remains a major clinical challenge. The present study identifies SCD1-mediated lipid desaturation as a metabolic vulnerability that can be exploited to increase the sensitivity of glioma cells to TMZ-induced cytotoxicity.Inhibition of SCD1 disrupts the SFA/MUFA balance, leading to the accumulation of saturated lipid species, induction of ER stress, and impairment of cellular stress-adaptation mechanisms. Our results indicate that these metabolic disturbances create a cellular environment that is less capable of tolerating the DNA damage and oxidative stress induced by TMZ. Consequently, combined SCD1 inhibition and TMZ treatment produced a marked reduction in cell viability and clonogenic growth compared with either treatment alone.

A particularly notable finding is the heightened sensitivity of IDH-mutant glioma cells to the combination therapy. IDH mutations are known to reprogram cellular metabolism and alter lipid composition, resulting in unique dependencies that distinguish these tumors from their IDH-wild-type counterparts. Previous work from our group demonstrated that IDH-mutant glioma cells preferentially incorporate MUFAs into ER and Golgi membranes, leading to organelle dysfunction and altered membrane architecture. The present findings extend this model by showing that inhibition of MUFA synthesis further compromises the ability of IDH-mutant cells to maintain membrane integrity and adapt to metabolic stress. As a result, these cells appear particularly vulnerable to additional insults such as TMZ-induced DNA damage.

Transcriptomic analyses revealed that the enhanced cytotoxicity of the combination treatment is associated with activation of multiple regulated cell death pathways. Signatures of both ferroptosis and necroptosis were detected following treatment with MF-438 and TMZ, suggesting that the therapeutic response cannot be attributed to a single mechanism of cell death. The activation of ferroptosis is consistent with our observations that SCD1 inhibition disrupts iron homeostasis, alters lipid composition, and weakens antioxidant defenses. At the same time, induction of necroptotic pathways indicates that severe metabolic and genotoxic stress may trigger parallel cell death programs, collectively amplifying tumor cell killing. The simultaneous engagement of multiple regulated death pathways may be particularly advantageous therapeutically, as it could limit the emergence of resistance mechanisms that commonly arise when only a single cell death pathway is targeted.

Another important observation was the induction of intracellular lipid transport pathways and neutral lipid metabolic processes following combination treatment. These transcriptional changes likely represent compensatory attempts by glioma cells to buffer excess saturated fatty acids and restore membrane homeostasis. However, these adaptive responses appear insufficient to overcome the profound metabolic stress imposed by combined SCD1 inhibition and TMZ treatment. The concurrent activation of the integrated ER stress response further supports the notion that disruption of lipid desaturation overwhelms the cellular machinery responsible for maintaining membrane and protein homeostasis.

Taken together, our results support a model in which SCD1 inhibition lowers the threshold for TMZ-induced cell death by simultaneously disrupting lipid metabolism, iron homeostasis, and redox balance. These alterations compromise the ability of glioma cells to repair damage and adapt to stress, ultimately promoting ferroptotic and necroptotic cell death. Importantly, the enhanced sensitivity observed in IDH-mutant glioma cells suggests that metabolic targeting of SCD1 may provide a therapeutic strategy that exploits vulnerabilities created by IDH1 mutant-driven metabolism.

## Conflict of Interest Statement

The authors declare no competing financial interests.

## Ethics Statement

All animal experiments were performed in accordance with institutional guidelines and were approved by NCI-Animal Use and Care Committee (ACUC), protocol number NOB-008.Transcriptomic and clinical data were obtained from publicly available, de-identified databases, including TCGA, CGGA, and GTEx (accessed through GEPIA3 and Gliovis platforms). No new human subjects were enrolled in this study, and no identifiable patient information was used.

## Funding Statement

This research was supported by the Intramural Research Program of the National Institutes of Health (NIH): ZIA BC011707-09 “The effect of therapeutics on mutated IDH1 cell lines and mice tumor models”. The contributions of the NIH author(s) are considered Works of the United States Government. This project has been funded in whole or in part with Federal funds from the National Cancer Institute, National Institutes of Health, Department of Health and Human Services, under Contract No. 75N91019D00024. The content of this publication does not necessarily reflect the views or policies of the Department of Health and Human Services, nor does mention of trade names, commercial products, or organizations imply endorsement by the U.S. Government.

## Data Availability Statement

All datasets generated during this study are available from the corresponding author upon reasonable request.

## Supplementary Materials

**Supplemental Figure S1.**
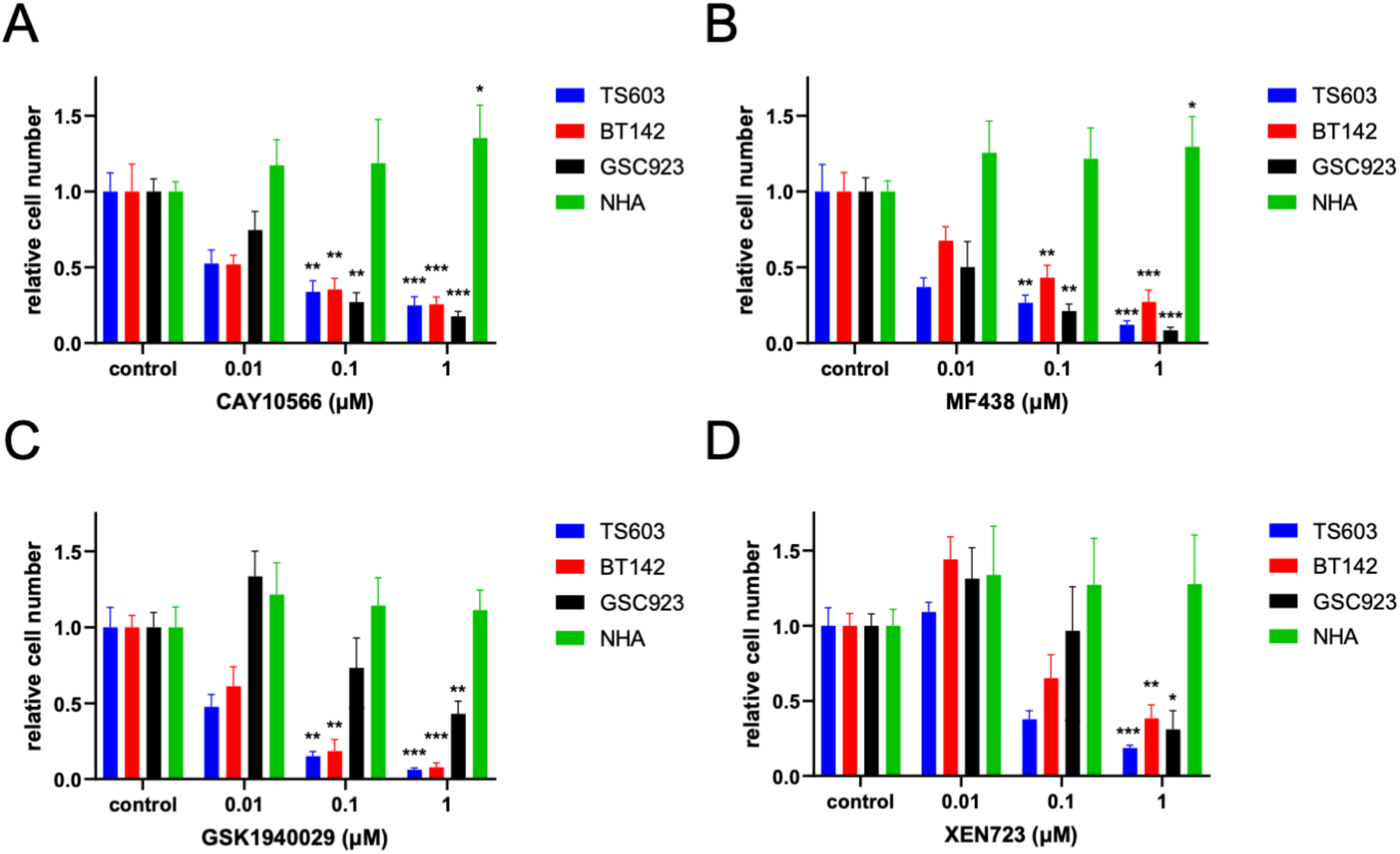
Normal Human astrocytes NHA cells are less affected by SCD inhibitor than glioma cells. The cell number of IDH1^mut^ cell lines (TS603, blue; BT142, red;), IDH^WT^ cell line (GSC923, black) and normal human astrocytes (NHA, green) was measured using trypan blue assay as a function of different SCD1 inhibitors: A: results for CAY10566; B: Results for MF438, C: Results for GSK1940029, D: Results for XEN723. All bar graphs are the average of at least three independent experiments (n=3) and the statistics were obtained using student t-test.

**Supplemental Figure S2.**
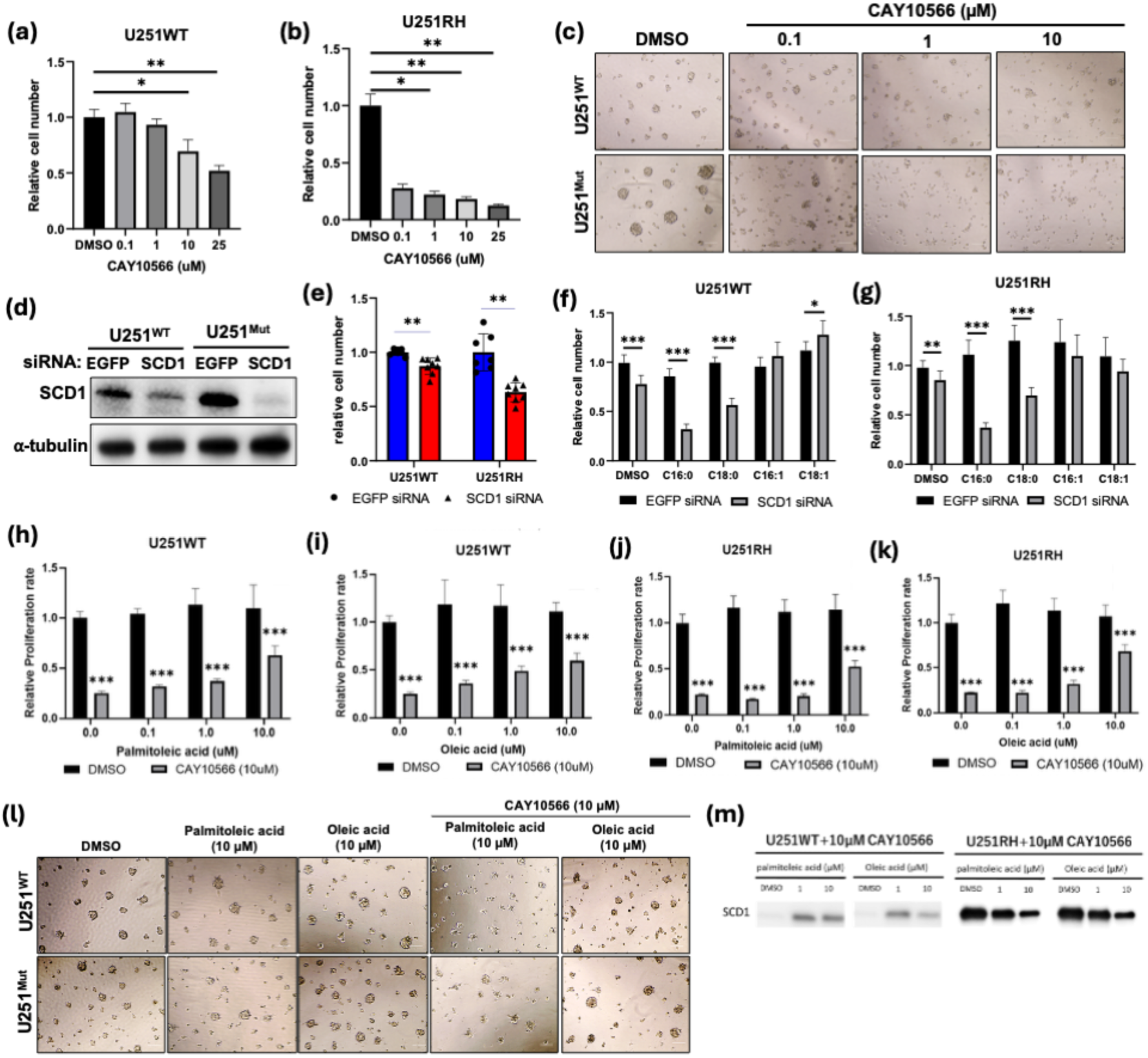
SCD1 inhibition or genetic knockdown affects the cell number and can be enhanced by SFA and rescued by MUFA addition. a), b) and c) IDH1WT and IDH1MUT cells treated with SCD2 inhibitor at increasing concentrations show decrease in cell proliferation. d) and e). Genetic knockdown of SCD1 via siRNA shows decreased proliferation more pronounced in IDH1^MUT^ cells compared with IDH1^WT^ ones. f) and g) Exogenous addition of SFA (C16:0 and C18:0) in induces further death in knockdown SCD1 cells siRNA in both IDH1^WT^ and IDH1^MUT^ cells while the addition of MUFAs (C16:1 and C18:1), rescues the relative cell number. h)-k). Addition of MUFA in dose-dependent manner rescues the relative cell number for both IDH1^WT^ and IDH1^MUT^ cells that were pretreated with SCD1 inhibitor. l) Neurophere images of SCD1 inhibitor treatment and the rescue with MUFAs for both cell lines. m) Western Blot analysis of SCD1 protein shows increased in expression with SCD1 inhibitor and decreased with the addition of MUFAs.

**Supplemental Figure S3.**
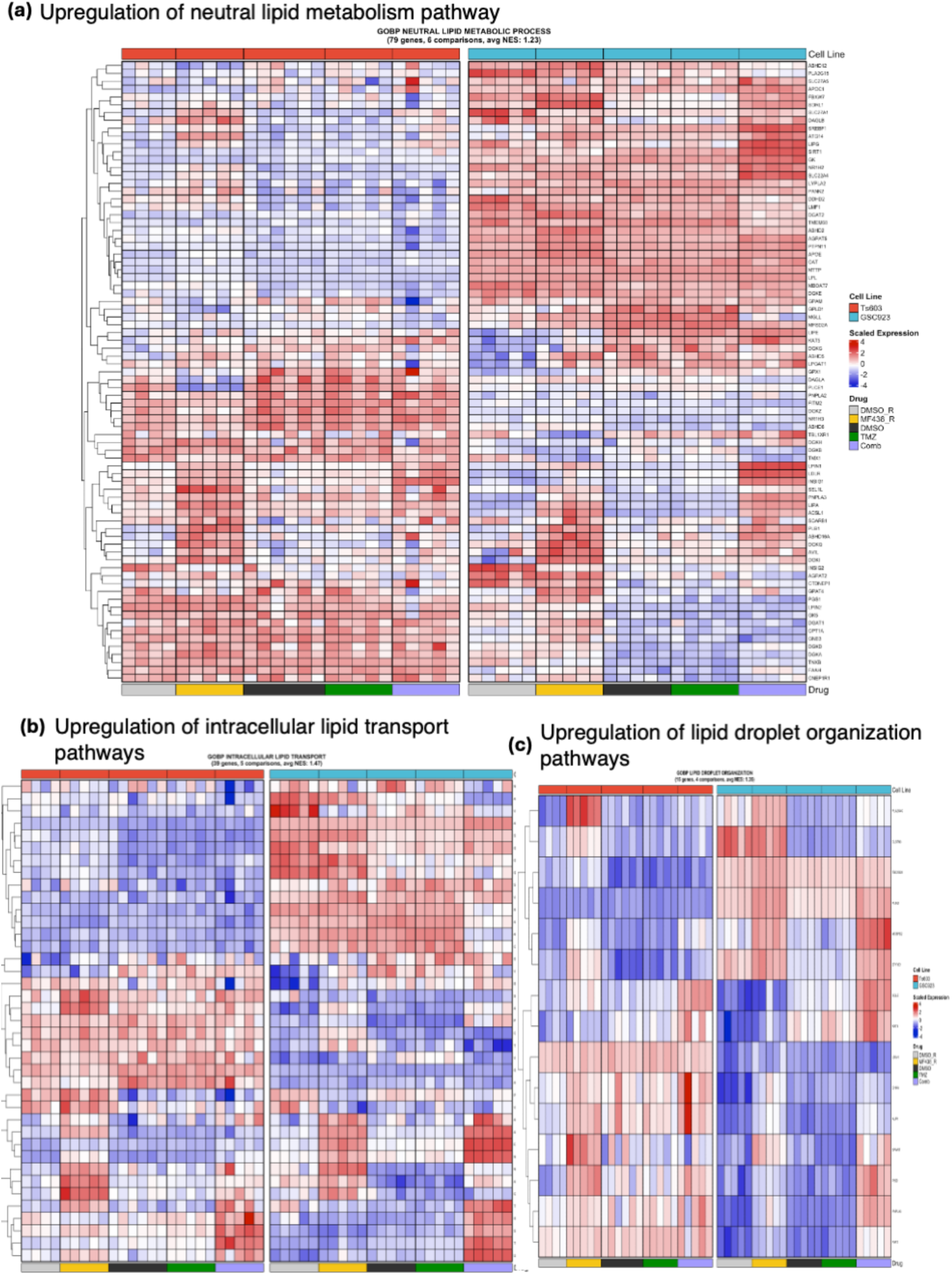
Combination treatment of MF-438 and TMZ upregulated lipid transport and metabolic pathways. **a**) RNA sequencing analysis of cell treated with MF-438, TMZ or combination show upregulation of lipid metabolic pathway more pronounced in GSC923. b) and c) RNA sequencing analysis of cell treated with MF-438, TMZ or combination show upregulation of lipid transport and lipid droplet organization.

**Supplemental Figure S4.**
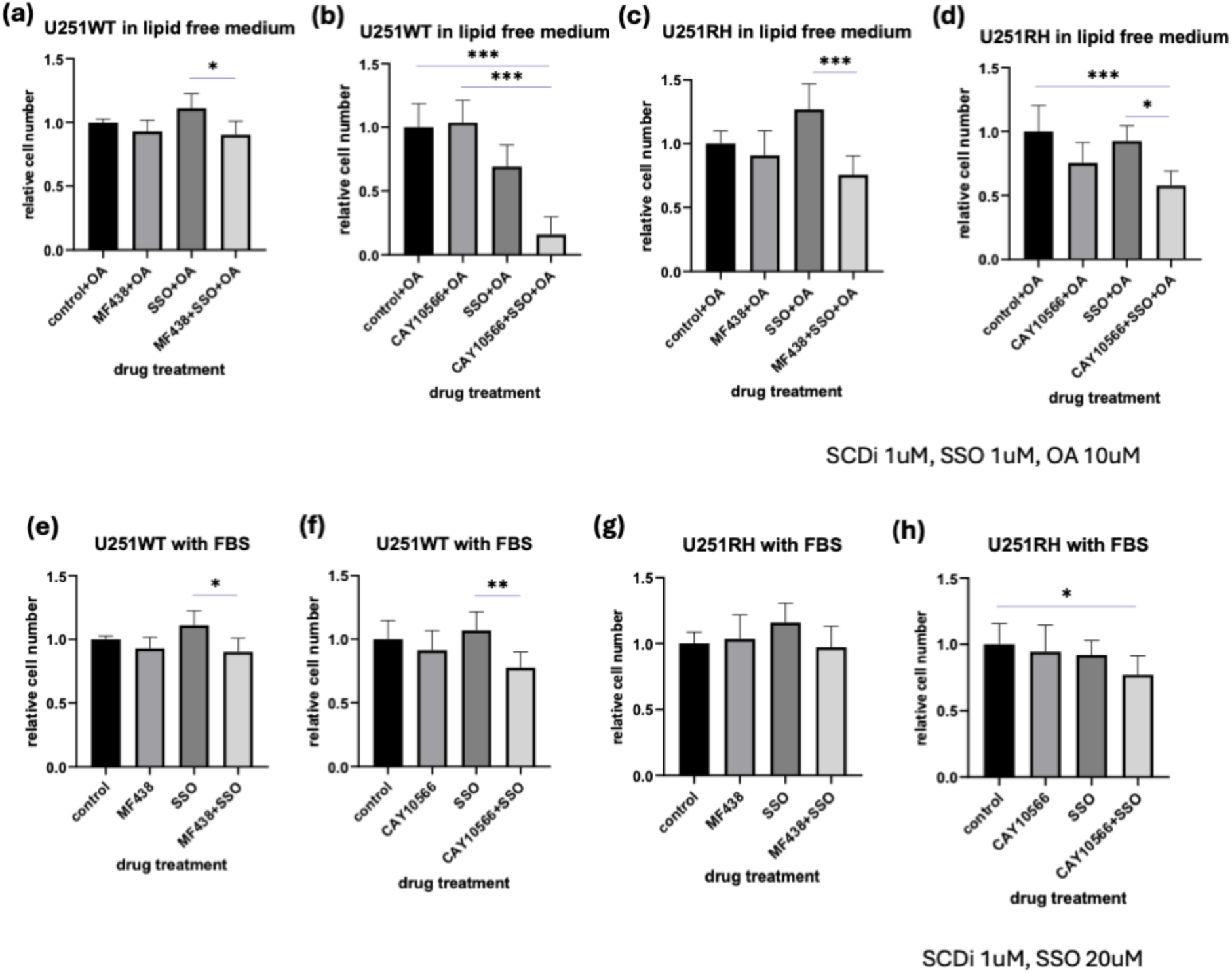
SCD1 Inhibition Combined with CD36 Blockade Synergistically Suppresses Glioma Proliferation by Targeting Exogenous MUFA Uptake. a)-d). The combination therapy between SCD inhibitor and CD36 inhibitor synergistically suppress glioma proliferation in exogenous added MUFA condition. e)-h). The combination therapy between SCD inhibitor and CD36 inhibitor synergistically suppress glioma proliferation in the medium with FBS.

## Notes

### Competing Interest Statement

The authors have declared no competing interest.

